# Sustaining Scholarly Infrastructures through Collective Action: The lessons that Olson can teach us

**DOI:** 10.1101/116756

**Authors:** Cameron Neylon

## Abstract

Infrastructures for data, such as repositories, curation systems, aggregators, indexes and standards are public goods. This means that finding sustainable economic models to support them is a challenge. This is due to free-loading, where someone who does not contribute to the support of the infrastructure nonetheless gains the benefit of it. The work of Mancur Olson (1974) suggests there are only three ways to address this for large groups: compulsion (often as some form of taxation) to support the infrastructure; the provision of non-collective (club) goods limited to those who contribute as a side-effect of providing the collective good; or mechanisms that lower the effective number of participants in the negotiation (oligopoly).

In this paper I use Olson’s framework to analyze existing scholarly infrastructures and proposals for the sustainability of new infrastructures. I argue that the focus on sustainability models prior to seeking a set of agreed governance principles is the wrong approach. Rather we need to understand how to navigate from club-like to public-like goods. We need to define the communities that contribute and identify club-like benefits for those contributors. We need interoperable principles of governance and resourcing to provide public-like goods and we should draw on the political economics of taxation to develop this.

## Introduction

### The provisioning problem for public scholarly goods

The fundamental political and economic challenge for groups is the provision of “public goods” or “general utility”. These are goods that are non-rivalrous – they can be infinitely shared – and non-excludable – it is difficult or impossible to stop someone using them. Classical economics tells us there is a provisioning problem for such goods, the rational individual actor will never contribute because whether they do or not, they can still benefit.

Infrastructures, such as repositories that provide access to data, articles and code, are very close to the ideal of public goods. These infrastructures require resources to build, resources to create the objects that they house, and resources to maintain over time. The public-good nature lies in a combination of the content itself, access to it, and the metadata that supports discovery and use of the objects they house.

Mancur Olson (1974) in *The Logic of Collective Action* developed a model of how such public goods can be provided, the circumstances under which they will not, and possible solutions to the dilemma. In particular, he discusses how group size has a profound influence on the provision of public goods, noting that optimal provision is only possible for small groups, or where the public good is a byproduct of the provision of non-public goods that are provided to contributors. Olson describes three different size ranges of groups. Those that are small enough to reach an agreement to provide the collective benefit, those that are too large, and those that lie in the middle where the collective good may be provided but at a level which is below optimal.

Olson’s description of the groups that can and cannot provide public goods maps closely onto scholarly infrastructures. Small communities frequently develop local infrastructures out of existing resources (and the contributions to these are usually biased strongly to the larger players in that community as Olson predicts). Large scale infrastructures are also provided by collaboration between small communities but in this case a community of funders (such as those that fund Europe Pubmed Central) or national governments (as is the case for physical infrastructures that are generally formed as Inter-governmental Organisation like CERN). The transition from small to large is challenging and “medium” sized infrastructures struggle to survive, moving from grant to grant, and in many cases shifting to a subscription model.

In the case of digital infrastructures, a public good (such as an online article or dataset) can be converted to a club good (made excludable) by placing an authentication barrier around it to restrict access to subscribers (as is the case for online subscription journals and databases). Buchannan and those that further developed his 1965 paper on the economics of clubs have probed how club goods and club size relate (Buchanan, 1965). A core finding is that such sustainable clubs have an equilibrium size that depends on congestion in access to good the extent to which it is purely non-rivalrous) and the value it provides. With digital resources congestion is low – although not zero – and the club can therefore presumably grow large. This in turn creates a challenge. Digital resources are not natively excludable; a technical barrier has to be put in place. As the group size rises the likelihood of “leakage” – sharing, or piracy (Lawson, 2017) if you prefer – increases. Thus resources are expended on strengthening excludability which leads to both economic and political costs as seen for example in the Open Access debate.

## Solutions to the provisioning problem

If our political goal is to provide large scale access, to make the goods created more public-like through the provision of shared infrastructures then we need to develop a political economics of this “public-making”. Olson provides three routes to creating sustainable infrastructures providing public goods:

Compulsion: The good is provided through a mechanism that requires contributions from the whole community. Closed Union Shops, where all workers in a given company are required to be members of a union are an example that Olson discusses in detail. Taxation is another example. In the scholarly infrastructure space, overhead and indirect costs taken by institutions are an example, as is the top slicing of funder budgets to provide infrastructures and services.
By-product: The public good is provided through a mechanism that additionally provides clublike and/or private goods to contributors. Olson discusses insurance schemes only available to members of mutual benefit societies that also lobby on behalf of their members. In the research enterprise publishers have to join Crossref to be able to assign Crossref DOIs to outputs (a club good). As a by-product the whole community has access to an interoperable and (largely) open metadata set with defined schemas and access points (a public good).
Effective oligopoly: There are far too many funders globally for them to (easily) agree a mechanism for all contributing to any shared scheme. However because a relatively small set of funders fund a substantial proportion of biomedical research they are able to agree mechanisms to fund data infrastructures such as Europe Pubmed Central. Other funders may contribute but many smaller ones will free-ride as Olson predicts.

The difficult truth that Olson articulates is that there is no mechanism that will directly lead to a large community supporting the provision of a large-scale public good infrastructure. Any successful sustainability model will depend on some mixture of these three approaches for resourcing. There are interesting models that quantitatively address these collective action problems. These include Assurance Contracts which are discussed by (Crow, 2013) as a means of addressing the size issues. Assurance contracts operate in a manner similar to many crow-funding approaches where a project only proceeds if sufficient contributions are made. These can be seen as ways for a community to implement compulsion on itself. Olson (1974) notes for instance that it was not uncommon for the majority of a group of workers to vote, even in private ballots, to form a closed union shop. This parallels an Assurance Contract.

If our challenge in delivering on the openness and transparency agenda is one of supporting the conversion of successful medium-scale club-like provision of infrastructure into open systems providing collective goods then we need to solve the political and economic problems of transitioning from the club state to a model that successfully provides a mix of these models.

## Case studies of scholarly infrastructures

In this section I examine a series of examples of scholarly infrastructures to given an outline of how Olson’s model can be applied for the research community. These case studies draw in part on statements made by data infrastructures for a session at the 2016 SciDataCon meeting that was part of the work of the OECD Expert Group on Data Infrastructure Sustainability, as well as some parallel examples drawn from the community. The examples are selected to show some parallels of history, variations in the models developed, as well as the attributes of the collective goods being provisioned.

### Cambridge Crystallographic Data Centre

The CCDC is a long standing not-for-profit that manages small molecule crystallographic data (Groom et al., 2016). The CCDC was founded over fifty years ago and is the dominant resource of its type in the chemical sciences. The CCDC has offered free online access to individual datasets for some time but access to various discovery and analytical systems requires a subscription (Bruno and Ward, 2016a, 2016b). Access to those benefits is a non-collective benefit of paying the subscription and a byproduct of providing the collective public-like good of a centralized data resource.

As the political and academic pressure for greater access has increased the Centre has opened up resources in a controlled way, but retaining key benefits for subscribers. A central argument for the model for CCDC is that the subscription/services model means that commercial interests provide a substantial amount of revenue. In turn the restrictions that protect this revenue create some – difficult to quantify – opportunity cost for the system as a whole.

Two options exist, shifting to a model for subscription (non-collective) benefits other than access to data resources, or an oligopoly model in which a consortium of interested industrial and public funders underwrite the costs of provision. While both of these are potentially plausible in principle the major challenge lies in the risks of managing a transition to a new model which cannot be guaranteed to work.

### The Arabidopsis Information Resource/Phoenix Bioinformatics

The Arabidopsis Information Resource provides plant genomic data resources to a wide community of biologists. In its original form as a US National Science Foundation funded project it offered these resources free online. However when the NSF progressively reduced and then removed funding the resource was obliged to find a new model (Reiser et al., 2016). It shifted from being a free resource to a limited access model with subscriptions for heavy users. In some senses it has travelled the opposite direction to CCDC to arrive at a similar point.

The current TAIR model is for metered access. Any IP address can access a number of pages free each month (currently 75). Heavy users will encounter a subscription page. As with CCDC subscribers gain additional non-collective benefits, in the case of TAIR unmetered access (Huala et al., 2016a). There is also a community building aspect as those who subscribe know they are contributing to the resource. TAIR is a long standing resource that provides labour-intensive curation and management of highly structured data. The availability of this data underpins the ability of plant scientists to be able to manipulate and study plant genomes and functionality and therefore provides a collective good.

The TAIR team (Huala et al., 2016b;Reiser et al., 2016) describe a high acceptance by the target community of core users of the new model. This may be close to an optimal case of how large a community can grow but still find a means of agreeing to fund a collective good. In particular, TAIR was a long standing resource, and had strong support from a relatively homogeneous community of disciplinary researchers. In addition, the crisis created by the loss of NSF funding created excellent conditions for the community to come together and solve a clear collective, and now internal, problem.

TAIR seeks to balance the need for revenue generation, and therefore non-collective benefits to members (i.e. database access), with public good provision through allowing limited access to light users. The potential opportunity cost is difficult to quantify. TAIR notes that actual usage has not reduced substantially since subscriptions were introduced. Nonetheless the model means that TAIR is more focused on member provisions and growing usage and membership than in developing the public good aspects of the resource.

> A pay for access model provides a business incentive to increase usage. Result: data curation, discovery and reuse are priorities, as they drive up revenue.
>
> — (Huala et al., 2016a)

The focus here is on membership provision, again emphasizing the (relatively small) community and identifying the means of showing the importance of their collective action in funding the resource. Both CCDC and TAIR operate in ways that are club-like in the sense of Buchannan’s club economics models. In both cases, the challenge of making the resources more like public goods than they currently are is less one of identifying plausible models than in underwriting an attempted transition to a system that cannot be guaranteed to work in practice.

### World-wide Protein Data Bank

The World-wide Protein Data Bank (wwPDB) is the global host for structural data on biomolecules. Its history is substantially different to that of the CCDC having been built from the beginning as an open resource (Berman, 2008;Berman et al., 2012). The reasons for the difference are both historical and cultural, and beyond the scope of this paper, but the history and expectations of the community are an important aspect of the differences (Byrd et al., 2016).

The PDB also experienced funding uncertainty in the early 2000s which was part of the motivation for the formation of the world-wide consortium in 2003. The international consortium is made up of members that seek funding from regional sources. In each region (the US, Europe, and Japan) a small set of funders support a local data centre (Byrd et al., 2016). This is effectively an oligopoly model for each region which is then managed across the regions in a way which provides some resilience if a single region were to fail.

The wwPDB is such a widely used and important resource that it is too crucial to be allowed to fail at this point. A small number of resources reach this level of importance but the PDB appears to be unique in being sustained through a network of regional grants. Other data resources in this class are funded through unusual direct arrangements (such as Genbank, Pubmed Central and other critical resources housed at the National Centre for Biological Informatics within the US National Library of Medicine) or through intergovernmental agreements such as the biological resources supported at the European Bioinformatics Institute.

The models for all these critical resources can be characterized as a mixture of compulsory taxation models (the resources are top-sliced from funder budgets such as NCBI) or oligopoly models (such as the intergovernmental agreement which funds the EBI). The intergovernmental oligopoly model is also the main model for large scale physical scholarly infrastructures such as CERN, the Institut Laue-Langevin and European Synchrotron Radiation Facility.

### Europe Pubmed Central

Europe Pubmed Central is a collaboration of European funders to provide a central resource for bibliographic information and full text resources of scholarly articles (PMC, 2015). It started as a collaboration of UK funders, with the Wellcome Trust and Medical Research Council as main drivers. It now counts 27 funders as members of the funding consortium (Europe Pubmed Central, 2016).

EuPMC is a clear example of an oligopology model. It started with the unilateral action of a small set of funders and has grown over time but the Wellcome Trust remains as the lead partner and contributions are based on the size of the contributing funders. A five year grant was awarded to the EBI to fund its continuation through to 2020. This involved some shift from the previous model which had been a collaboration across multiple sites. While not strictly a tendering model this shows the principle that the funders could decide to continue the resource by funding another party to run it. This emphasizes the funder and therefore oligopoly-directed nature of the funding and provision.

It is difficult to see how EuPMC could transition to a different sustainability model. It could be conceivable to expand the membership base to include institutions or publishers in some form but the benefits of this are unclear. The public good provision provides some non-collective benefits to contributing funders in the form of a designated repository that their grantees can be required to use (Europe Pubmed Central, n.d.), however the main goal behind its funding is to shift the research enterprise towards an open access model, a clear case of a small group deciding to directly fund a collective good.

### Crossref (and ORCID)

Crossref provides a collective public-like good in the form of freely accessible bibliographic metadata and the infrastructure that supports it. Financially Crossref is supported by member publishers who gain the ability to assign Crossref Digital Object Identifiers (DOIs) to work that they publish alongside access to a set of services that aid in metadata creation, discovery, and validation of content (e.g. for plagiarism through the Similarity Check service).

Crossref was started essentially by fiat by a small number of publishers. In practice between five and nine publishers dominate the market (Lariviére et al., 2015). The setup and early funding of Crossref, while more complex than a simple agreement (Crossref, 2009), was successful largely because a group representing a substantial proportion of scholarly publishing could come together. This is a classic oligopoly.

Crossref offered a non-collective membership benefits from its inception, including the ability to assign DOIs and the traffic that results from referral. A report by Greenhouse Associates in 2004 for an unnamed publisher noted the direct financial benefits that were arising for publisher members engaging with the Crossref system (Greenhouse Associates, 2004). These benefits can be framed as a byproduct of producing the public good of a shared metadata system.

Finally today, membership of Crossref is effectively compulsory for any serious publisher of scholarly articles in STEM subjects, and increasingly for humanities and social sciences as well as scholarly book publishers. It is just part of the costs of doing business for a publisher. A compulsory, tax-like, part of the system.

It is helpful to contrast the growth of ORCID (Haak et al., 2012). ORCID, the provider of identifiers for contributors to scholarly work. Publishers were a core of the early group of advocates, alongside a few funders. It was publishers that provided the majority of the initial financing. Over several years funders started to contribute. A key point is that while a small group of publishers dominate the market, the group of important funders is larger. Olson would therefore predict that publishers would agree collective action before funders.

Universities and research institutions have been very slow to engage with ORCID despite having the potential in many ways to reap the most benefits. However the most important benefits will only arise when there is substantial adoption. Here we see the classic adoption problem described by Olson. There are too many universities and research institutions, even if we restrict ourselves to the traditional Euro-American centres of scholarship to easily solve the collective action problem. Some progress has been made over the past few years, howeverthis has been primarily due to national coordination actions (Paglione, 2016). Coordination at a national level poses less of a collective action problem than across thousands of institutions. An important lesson from the ORCID experience is to seek to shift the challenge of coordination to a level where the number of players is small enough to solve the collective action problem.

### AddGene

Addgene is an interesting counterpoint to the other resources here because it provides access to a physical resource. Addgene is a not-for-profit organization founded in 2004 to provide access to plasmids, DNA materials that are used as tools and resources across many areas of biology. Addgene will archive store and commit to distributing plasmids from researchers for free (Kamens, 2015). To obtain a plasmid users need to pay a fee (Addgene, n.d.).

The Addgene model works because it couples a collective benefit (central collection, archiving, and quality assurance of valuable materials) with a non-collective benefit (access to a physical rivalrous material, the plasmid itself). For users paying the relatively modest fee to obtain a plasmid is well justified because Addgene can provide quality assurance. While these materials circulate widely between labs and often without any charges, the cost of lost time if the fragile materials are found to have been degraded in transit can easily outweigh the apparent upfront cost of obtaining the quality assured material from Addgene. That saving in time, at the cost of an upfront fee, is a non-collective benefit. This is Olson’s by-product model.

By centralizing the availability and management of plasmids Addgene creates a central information resource that is more valuable than the sum of its parts. The quality assurance and archiving, even the distribution processes are much cheaper when managed at scale and Addgene is effective at exploiting that scale to create value for those paying for the resource.

## The politics of compulsion and the need for shared governance

Of the three approaches to sustainability, it is generally the second which infrastructures are expected to pursue as they grow. Membership models can work in those cases where there are club goods being created which attract members. Training experiences or access to valued meetings are possible examples. In the wider world this parallels the "Patreon" model where members get exclusive access to some materials, access to a person (or more generally expertise), or a say in setting priorities. Much of this mirrors the roles that Scholarly Societies play or at least could play.

In the scholarly infrastructures space the compulsion/taxation and oligopoly approach are very similar in practice as top slicing by a group of funders amounts to a tax on overall research funds. Some membership models also approach the level of compulsion. While this is rare in scholarly communities, it is common in professional communities such as medicine, law, and some areas of engineering. Schemes offering professional certification (including the validation of degree programs by scholarly societies) blur this boundary as well.

The word “compulsion” is pejorative but there are many activities within the work of researchers that are compulsory. Gaining a doctorate, publishing at some level, having access to the scholarly literature in some form are all effectively compulsory. These forms of compulsion (or call them “social expectations” if you prefer) are considered acceptable because the fit within a known and understood system of rules. Systems of taxations are acceptable, according to Adam Smith (Smith, 1776) where there is proportionality, predictability, convenience and efficiency. Today we would probably add representation in governance and sustainability. This requires us to build institutions, in the sense that Elinor Ostrom describes them: “institutions are the prescriptions that humans use to organize all forms of repetitive and structured interactions” (Ostrom, 2005). Much of political economics is bound up in trying to justify post-hoc the provision of institutions like governments, courts, the law by inventing things like “the social contract”. Our advantage as a scholarly community, or communities, is that we can explicitly develop and agree principles of operation as a way of reducing the costs of creating institutions.

### A common set of principles for foundational infrastructures

Building institutions is hard. It takes resources. To reduce costs, it makes sense to build templates; sets of agreed principles under which such institutions and systems should operate. If our communities can sign up to a set of principles up front, then building institutions and infrastructures that reflect those principles should become a lot easier.

To address this a draft set of principles were developed to provoke a conversation about the governance and management of these infrastructures (Bilder et al., 2015). These principles rest on three pillars: transparency and community governance; financial sustainability, efficiency and commitment to community needs; and mechanisms to protect integrity and manage and mitigate the risk of failures. They draw from the observation of successes in providing foundational infrastructures and seek to generalise these. The focus is on building *trustworthy* institutions.

The principles also map quite closely to Smiths’ four principles for sound taxation. The commitment to representation is more modern, but a core part of building stable community and trust. The concept of enabling a community to fork a project through committing to Open Source and Open Data – while modern in its approach – could be framed an expression of the principles of efficiency and predictability. The intent of the Infrastructure Principles is to ensure that the community, or club, has a series of mechanisms that provide consistency and predictability. The first line is community governance and control, the second is financial stability. Forkability ensures there is a reliable path to deal with a worst case scenario in which an infrastructure is co-opted or “goes rogue”.

Smith’s principles and the Infrastructure Principles of Bilder et al. are examples of pattern languages in the sense advanced by Bollier and Helfrich (Bollier and Helfrich, 2015; Helfrich, 2015). They also speak to a broader goal of creating good culture that supports infrastructures. Ultimately we will need a broader understanding of how institutions and culture evolve over time as the communities they relate to scale (Hartley and Potts, 2014;Wilson et al., 2014). In the shorter term, the identification of patterns and an understanding of where they can be applied provides significant promise of making progress.

Such patterns (or a future refinement or replacement for them) can serve in two ways. First they could be used to set out the minimum requirements for governance and sustainability that are required before funders are willing to directly fund (the oligogopoly or tax mechanisms). Second they provide a template for a developing club, either one built in a community sufficiently small to bootstrap its funding, or one that has found a byproduct model, to demonstrate its ability to make the transition from *club* to *infrastructure*.

## Predictions and practical consequences

While the case studies are helpful, the above remains an abstract argument. What are its practical consequences for actually sustaining scholarly infrastructures? First we can make a prediction to be tested: All sustainable scholarly infrastructures providing collective (public-like) goods to the research community will be funded by either one of the three identified models (taxation, byproduct, oligopoly) or some combination of them.

Second, we can look at stable long standing infrastructures (Crossref, Protein Data Bank, NCBI, arXiv, SSRN) and note that in most cases governance arrangements are an accident of history and were not explicitly planned. Crises of financial sustainability (or challenges of expansion) for these organisations are often coupled to or lead to a crisis in governance, and in some cases a breakdown of community trust. Changes are therefore often made to governance in response to a specific crisis.

Where there is governance planning it frequently adopts a best practice model which looks for successful examples to draw from. It is not often based on worse case scenario planning. This is a problem. We can learn as much from failures of sustainability and their relationship to governance arrangements as from successes.

We can look towards using these economic models to investigate what forms of funding and organization are likely to work for the future. An initial suggestion is that membership-fee based sustainability models will only work where a non-collective benefit is provided. In turn this means identifying where such non-collective benefits are a viable model. A deeper understanding of which non-collective benefits are appropriate will be valuable and will help to address the assumption that membership and subscription systems can only be tied to access to content. It will also be important to identify where non-collective benefits are not a viable model, and to avoid forcing the model on organisations for which it cannot work.

With this deeper understanding of how models work, and in particular how they relate to the scale of communities, it should be possible to create pathways to sustainability. These pathways would link the growth of infrastructures from club-like services, through community adoption, to the creation of public goods, with the appropriate sustainability models and financial instruments to support them, and a graduated set of governance requirements appropriate to the stage of development.

Previous work on sustaining infrastructures has mostly focused on examination of specific financial models (Bastow and Leonelli, 2010;Ember and Hammisch, 2013). Crow’s report for Knowledge Exchange offers the major exception here and its practical development of the options that Assurance Contracts may offer is an important complement to this paper (Crow, 2013). While the issue of financial sustainability is key, and the political discussion of the balance of resource allocation between platforms and new research is important, my claim here is that most studies have not addressed two important dimensions in sufficient detail.

Firstly that infrastructures need to be seen as both sustaining and being sustained by the communities that they serve. It is the *political economy* that needs to be addressed, not simply the economics. Secondly that the size of the community and the scale of the infrastructure is a critical aspect of defining what sustainability models can work, and that sustainability models must therefore change through the growth and development of an infrastructure.

Above all the key is to learn from our experiences, as well as from the theory of economics and governance to identify the patterns and templates that provide resilient organisations and infrastructures that through being trustworthy earn the trust of their communities through both the good times and the bad.

